## Supplementary Figures for "Glutamine deficiency in solid tumors confers resistance to ribosomal RNA synthesis inhibitors" (from another version of this preprint: version 1, 2021-12-04)

**Supplementary Fig. 1**

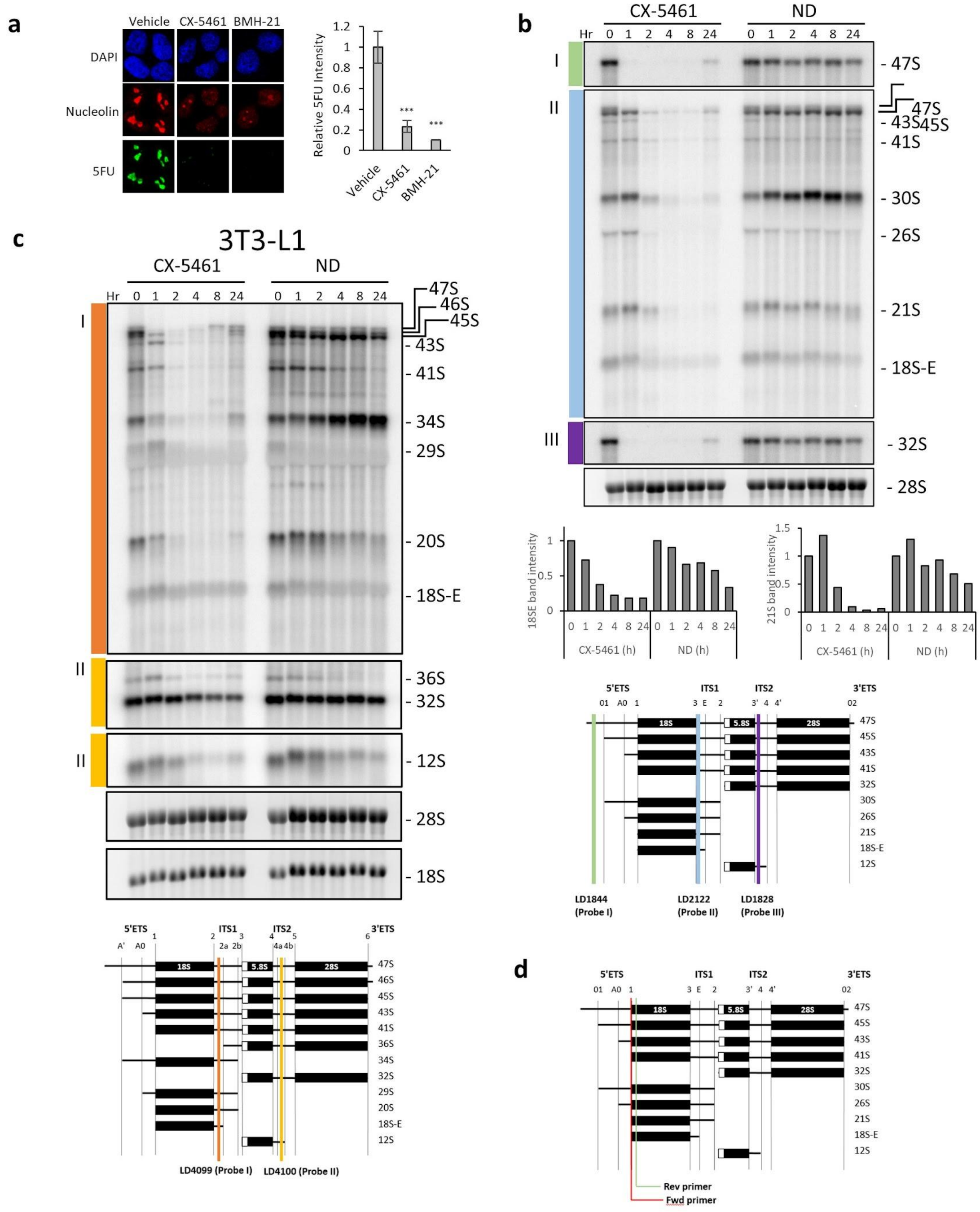


**Supplementary Fig. 1**

(a) CX-5461 and BMH-21 inhibits 5-Fluorouridine (5FU) uptake in HCT116 cells. HCT116 cells were treated to 10 µM CX-5461 or 1 µM BMH-21 for 1 hr, and nascent RNA was pulse-labeled with 5FU for 5 minutes.

(b) Top: Pre-rRNA expression in HCT116 cells treated to 10 µM CX-5461 or ND as assessed using northern blotting. Middle: Quantification of 21S and 18S-E species in response to CX-5461 or ND treatment. Bottom: binding sites of northern blot probes.

(c) Top: Pre-rRNA expression in mouse 3T3-L1 cells treated to 10 µM CX-5461 or ND as assessed using northern blotting. Bottom: binding sites of northern blot probes.

(d) Binding sites of the 5’ETS pre-rRNA qRT-PCR primers. The forward primer recognizes site 01 of the pre-rRNA, which is cleaved upon maturation. Mature rRNAs do not contain the site 01 sequence.

**Supplementary Fig. 2**

**a**

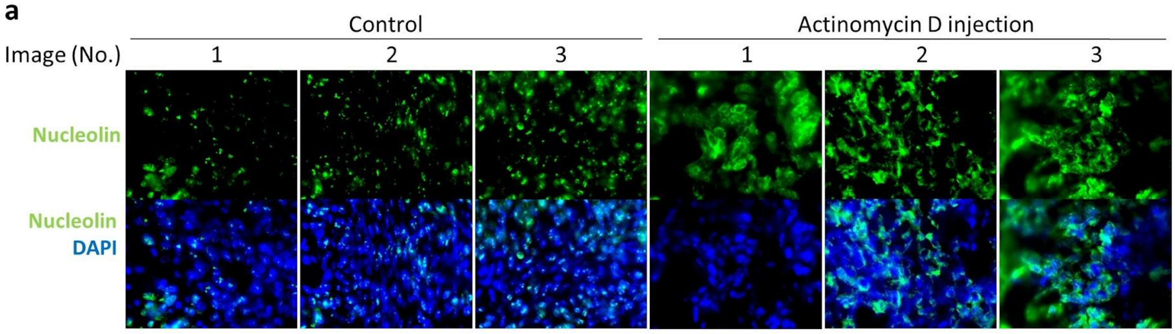


**b**

| Compound | Periphery (nmol/gram) | Core (nmol/gram) | Periphery (normalized to 1) | Core (relative to periphery) | p-value |
|---|---|---|---|---|---|
| Urocanate | 602.81 | 14.07 | 1.0 | 0.023 | 0.008991784 |
| Histamine | 195.42 | 20.17 | 1.0 | 0.103 | 0.026806533 |
| 5-Oxoproline | 937.97 | 138.45 | 1.0 | 0.148 | 0.01342651 |
| Arg | 751.84 | 139.44 | 1.0 | 0.185 | 0.0012917 |
| Cytidine | 9.86 | 2.69 | 1.0 | 0.273 | 0.010154887 |
| Glutathione(red) | 2533.56 | 744.07 | 1.0 | 0.294 | 0.000529169 |
| Ser | 2629.26 | 793.44 | 1.0 | 0.302 | 0.003870969 |
| Anserine | 7.06 | 2.33 | 1.0 | 0.330 | 0.032765351 |
| IMP | 563.22 | 188.46 | 1.0 | 0.335 | 0.014767429 |
| Argininosuccinate | 28.64 | 10.18 | 1.0 | 0.356 | 0.000503562 |
| UDP | 48.81 | 17.70 | 1.0 | 0.363 | 0.005865504 |
| Glycerophosphate | 238.82 | 88.59 | 1.0 | 0.371 | 0.000424676 |
| UMP | 579.36 | 218.28 | 1.0 | 0.377 | 0.00149572 |
| CDP | 11.04 | 4.43 | 1.0 | 0.401 | 0.001914604 |
| AMP | 609.28 | 259.65 | 1.0 | 0.426 | 0.011314459 |
| Carnosine | 8.65 | 3.74 | 1.0 | 0.432 | 0.000259768 |
| Cystathionine | 10.80 | 4.68 | 1.0 | 0.433 | 0.022993214 |
| CMP | 87.89 | 38.07 | 1.0 | 0.433 | 1.96385E-05 |
| UDP-glucose | 209.63 | 90.85 | 1.0 | 0.433 | 0.001532262 |
| GMP | 286.98 | 127.93 | 1.0 | 0.446 | 0.002287422 |
| Glycerophosphorylcholine | 61.40 | 27.61 | 1.0 | 0.450 | 0.002795165 |
| gamma-Glu-cys | 5.86 | 2.63 | 1.0 | 0.450 | 0.009944522 |
| UDP-glucuronate | 72.42 | 32.63 | 1.0 | 0.451 | 0.00030003 |
| Hypotaurine | 233.51 | 106.61 | 1.0 | 0.457 | 0.006931956 |
| GDP-mannose | 5.04 | 2.31 | 1.0 | 0.458 | 0.033945489 |
| Gln | 3152.53 | 1475.04 | 1.0 | 0.468 | 2.78503E-06 |
| Spermidine | 21.77 | 10.40 | 1.0 | 0.478 | 0.000682741 |
| UTP | 5.39 | 2.58 | 1.0 | 0.480 | 0.020690314 |
| UDP-N-acetylglucosamine | 479.11 | 236.26 | 1.0 | 0.493 | 0.000413227 |
| His | 311.53 | 159.12 | 1.0 | 0.511 | 0.005440344 |
| Val | 801.58 | 409.45 | 1.0 | 0.511 | 0.003877367 |
| dTMP | 9.64 | 4.94 | 1.0 | 0.512 | 0.001589071 |
| Trehalose 6-phosphate | 62.13 | 31.99 | 1.0 | 0.515 | 0.002041459 |
| Asn | 589.84 | 307.43 | 1.0 | 0.521 | 5.23679E-05 |
| Ru5P | 54.98 | 29.25 | 1.0 | 0.532 | 0.009278918 |
| NADH | 5.41 | 2.90 | 1.0 | 0.536 | 0.01348221 |
| Phe | 357.79 | 194.47 | 1.0 | 0.544 | 0.003501808 |
| SAM+ | 26.04 | 14.55 | 1.0 | 0.559 | 0.001199224 |
| ADP | 170.88 | 95.46 | 1.0 | 0.559 | 0.012560832 |
| beta-Ala | 138.47 | 79.41 | 1.0 | 0.573 | 0.000314044 |
| 5-Aminolevulinate | 1.54 | 0.89 | 1.0 | 0.575 | 0.030835195 |
| G1P | 22.63 | 13.03 | 1.0 | 0.576 | 0.013847607 |
| N8-Acetylspermidine | 6.69 | 3.87 | 1.0 | 0.578 | 0.025265653 |
| N-Acetylaspartate | 241.99 | 139.98 | 1.0 | 0.578 | 0.005764296 |
| CDP-choline | 45.01 | 26.08 | 1.0 | 0.579 | 0.000593962 |
| 2AB | 27.33 | 15.95 | 1.0 | 0.584 | 0.041846706 |
| Met | 235.94 | 137.88 | 1.0 | 0.584 | 0.003469603 |
| Tyr | 388.58 | 228.06 | 1.0 | 0.587 | 0.001016928 |
| CMP-N-acetylneuraminate | 16.36 | 9.75 | 1.0 | 0.596 | 0.001850367 |
| N1,N12-Diacetylspermine | 11.54 | 20.00 | 1.0 | 1.733 | 0.013976182 |

**Supplementary Fig. 2**

(a) Immunofluorescence staining of nucleolin in HCT116 tumors. HCT116 tumors were intratumorally injected with actinomycin D (1.33 mM) and were collected after 30 min for cryosectioning and immunofluorescence. Three representative images are shown per treatment.

(b) Metabolome analysis of HCT116 periphery and core tissues (n=10). Twenty metabolites (black text) were downregulated in core tissues, and three metabolites (red text) were upregulated.

**Supplementary Fig. 3**

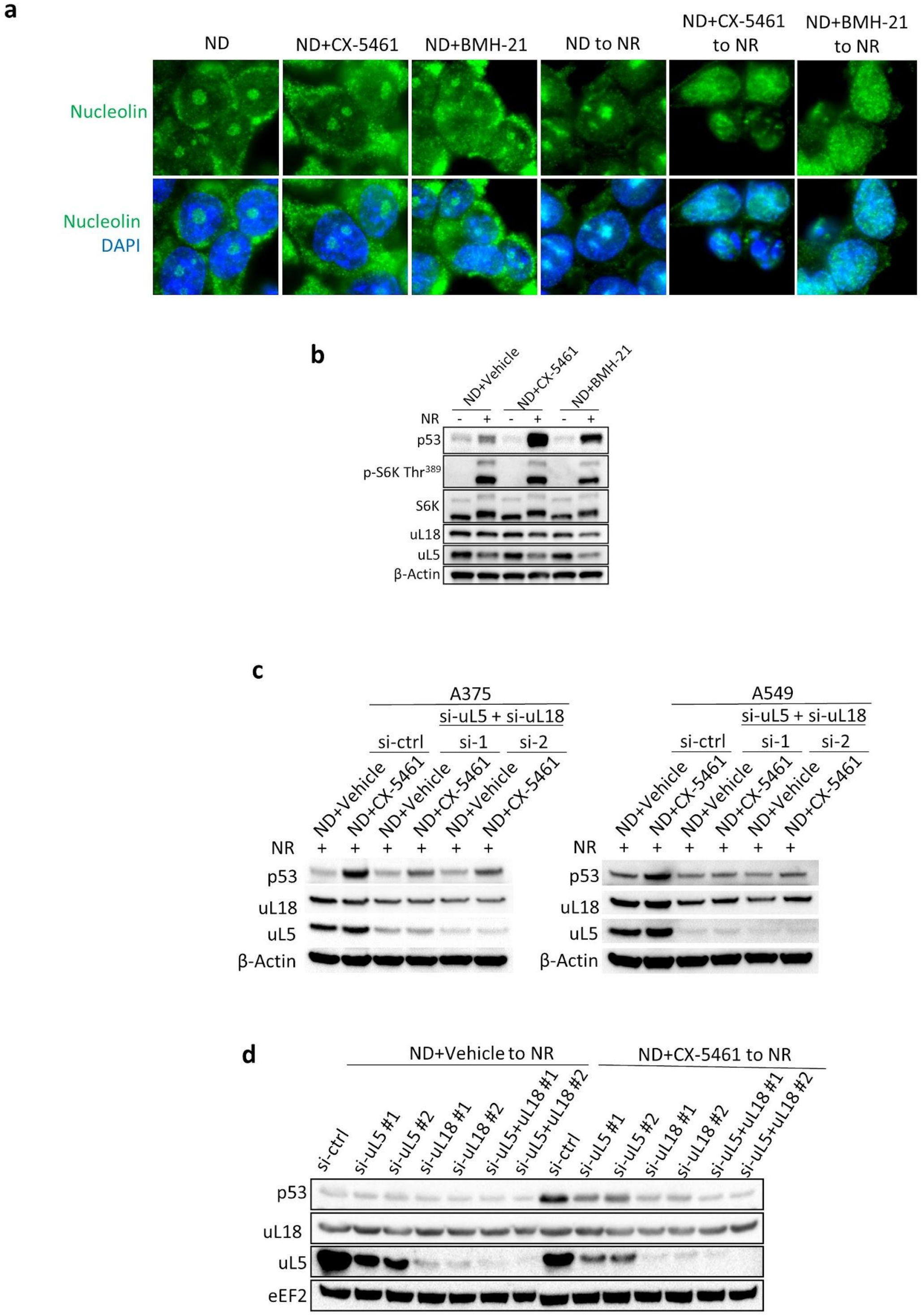


### Supplementary Fig. 3

(a) Nucleolin immunofluorescence on HCT116 cells treated to ND with vehicle ($NaH_2PO_4$), CX-5461 (10 µM), or BMH-21 (1 µM) for 24 h. After 24 h, cells were exposed to NR for 1 h.

(b) NR induces p53 in pre-rRNA depleted A375 cells. A375 cells were cultured under ND with vehicle ($NaH_2PO_4$), CX-5461 (10 µM), or BMH-21 (1 µM) for 24 h; after 24 h,

the treatment medium was replaced with a drug-free basal medium (NR) for 8 h.
(c) Co-knockdown of uL5 and uL18 by siRNA decreases NR-mediated p53 activation. A375 and A549 cells were transfected with control or two different sets of siRNAs targeting uL5 and uL18. After 48 h, the cells were placed under ND with CX-5461 (10 μM) for 24 h. After 24 h, the treatment medium was replaced with DMEM for 8 h and the protein lysate was collected for western blotting.
(d) Independently silencing uL5 or uL18 inhibits NR-mediated p53 activation. HCT116 cells were transfected with control or uL5/uL18 siRNAs. After 48 h, the cells were placed under ND with CX-5461 (10 μM) for 24 h. After 24 h, the treatment medium was replaced with DMEM for 8 h and the protein lysate was collected for western blotting.

**Supplementary Fig. 4**

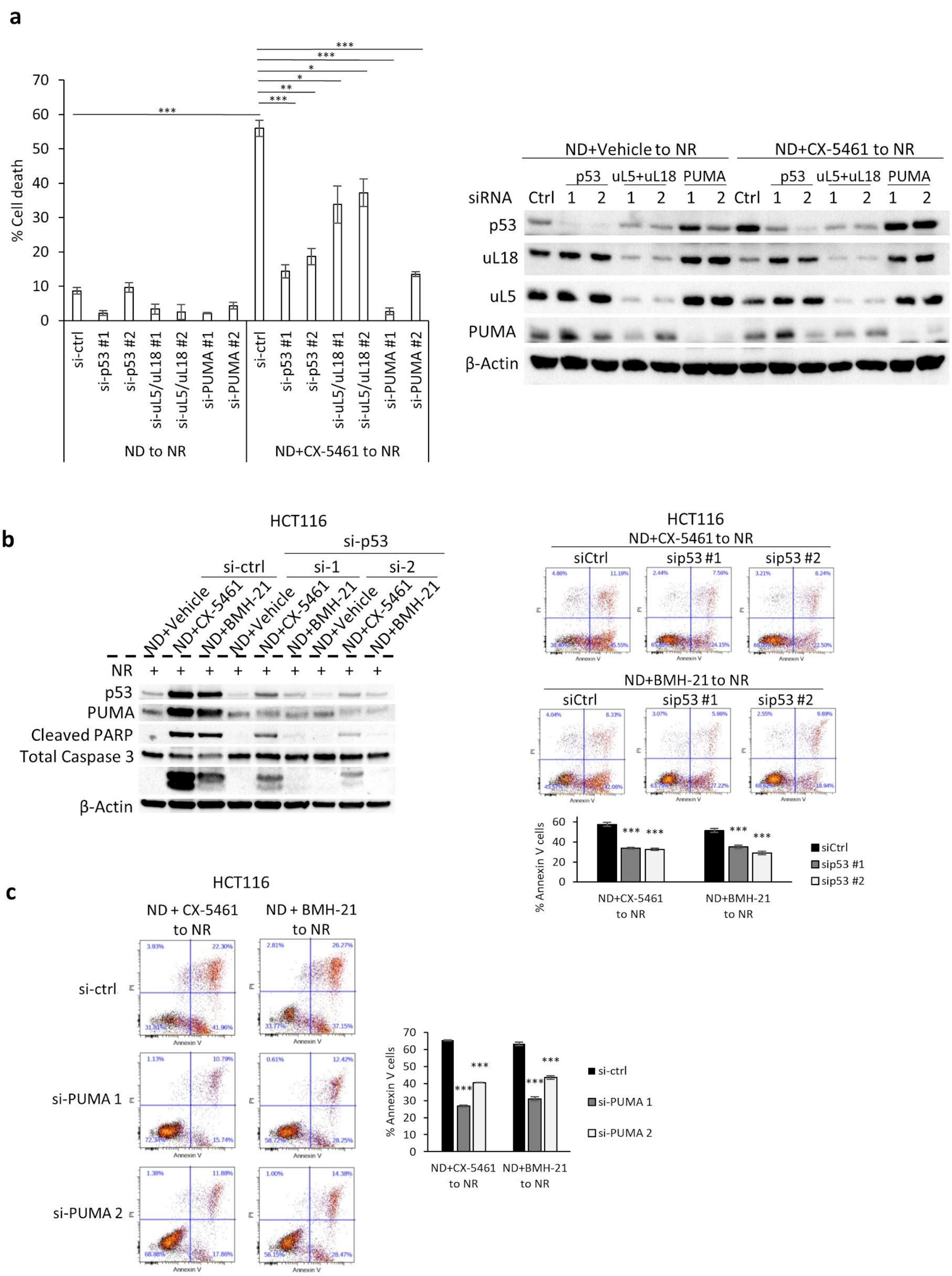


**Supplementary Fig. 4**

(a) Silencing uL5/uL18, p53, or PUMA, decreases NR-mediated cell death. HCT116 cells were transfected with siRNAs targeting uL5+uL18, p53, or PUMA. After 48 h, the cells were placed under ND with vehicle or CX-5461 (10 μM) for 24 h. After 24 h, the treatment medium was replaced with DMEM for 8 h and the protein lysate was collected for western blotting. Left: Percentage of cell death as assessed using the

trypan blue exclusion assay. Right: Expression of p53, uL18, uL5, and PUMA as assessed using western blotting.
(b) Silencing p53 decreases NR-mediated apoptosis in HCT116 cells. HCT116 were transfected with scramble or two independent p53 siRNAs. After 48 h, cells were placed under ND with vehicle ($NaH_2PO_4$), CX-5461 (10 μM), or BMH-21 (1 μM). After 24 h, the treatment medium was replaced with a drug-free basal medium for 24 h. Left: Expression of cleaved caspase 3 and PARP as assessed using western blotting. Right: Percentage of Annexin V+ cells as assessed using flow cytometry.
(c) PUMA siRNA knockdown inhibits NR-mediated apoptosis in pre-rRNA depleted HCT116 cells, as determined using Annexin V-PI flow cytometry. HCT116 were transfected with scramble or two independent PUMA siRNAs. After 48 h, cells were placed under ND with vehicle ($NaH_2PO_4$), CX-5461 (10 μM), or BMH-21 (1 μM). After 24 h, the treatment medium was replaced with a drug-free basal medium and Annexin V-PI flow cytometry was performed. Right: quantification of apoptosis (Annexin V+ cells) for the indicated conditions.

**Supplementary Fig. 5**

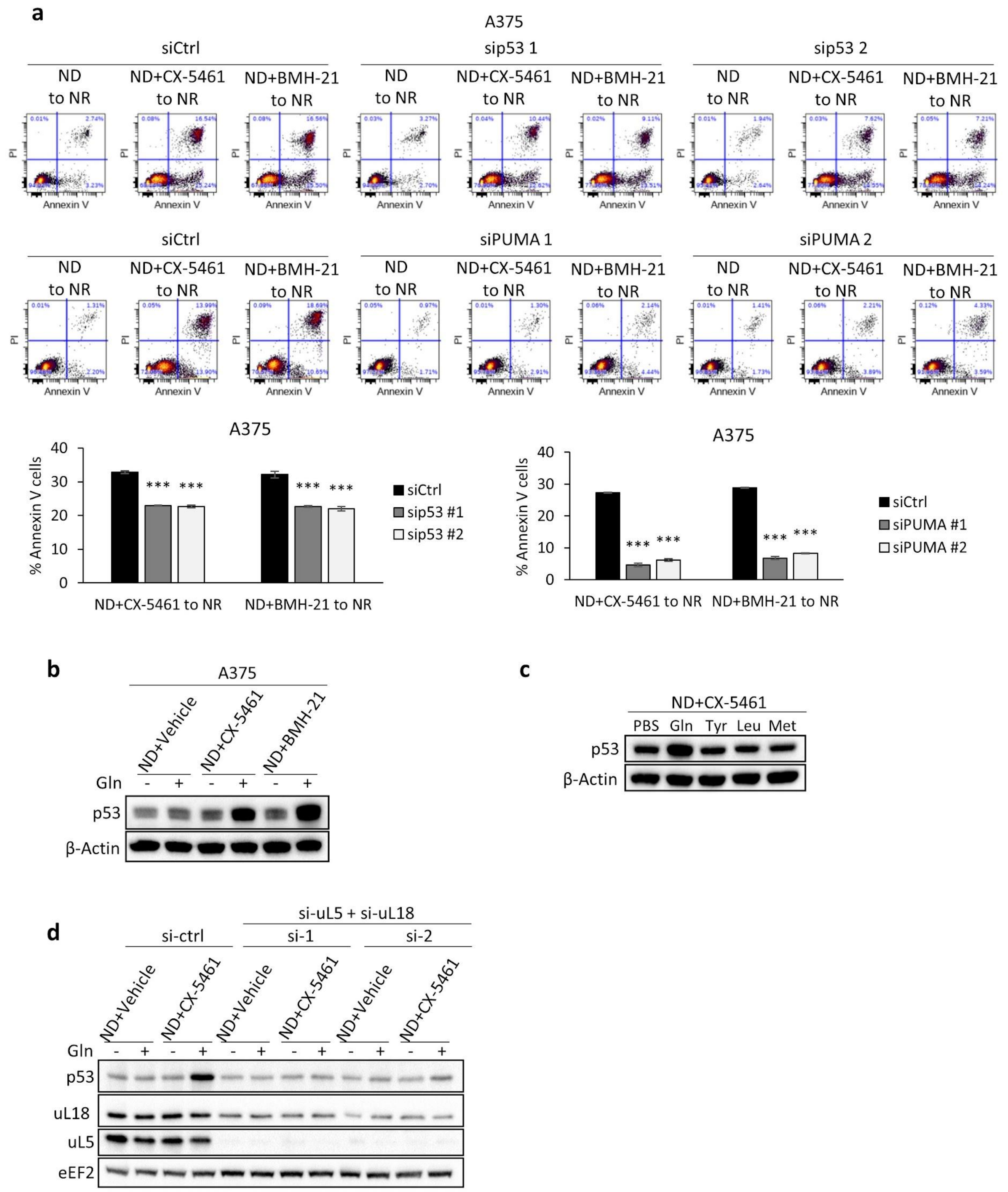


### Supplementary Fig. 5

(a) p53 or PUMA knockdown by siRNA inhibits NR-mediated apoptosis in A375 cells, as assessed by Annexin V-PI flow cytometry. A375 were transfected with siRNAs, and after 48 h, cells were placed under ND with vehicle ($NaH_2PO_4$), CX-5461 (10 µM), or BMH-21 (1 µM). After 24 h, the treatment medium was replaced with a drug-free basal medium and flow cytometry was performed after 24 h NR

(b) Glutamine (gln) activates p53 in pre-rRNA depleted A375 cells as assessed by western blotting. A375 were cultured in ND medium with vehicle ($NaH_2PO_4$),

CX-5461 (10 μM), or BMH-21 (1 μM) for 24 h. After 24 h the old treatment media was replaced with ND media containing PBS or Gln (4 mM) for 8 h.
(c) Glutamine (gln) but not tyrosine (tyr), leucine (leu), or methionine (met) activates p53 in pre-rRNA deficient HCT116 cells. HCT116 cells were placed under ND with CX-5461 (10 μM) for 24 h. After 24 h the old treatment media was replaced with ND media containing PBS, glutamine (4 mM), tyrosine (0.1 mg/mL), leucine (0.1 mg/mL), or methionine (0.03 mg/mL). The concentration of amino acids added back to the ND media is equal to the concentrations in the basal media.
(d) Dual siRNA knockdown of uL5 and uL18 inhibits glutamine (Gln)-mediated p53 activation. HCT116 cells were transfected with control or two different sets of siRNAs targeting RPL5 and RPL11. After 48 h, cells were placed under ND with CX-5461 (10 μM) for 24 h. After 24 h the old treatment media was replaced with ND media containing PBS or Gln (4 mM) for 8 h.

**Supplementary Fig. 6**

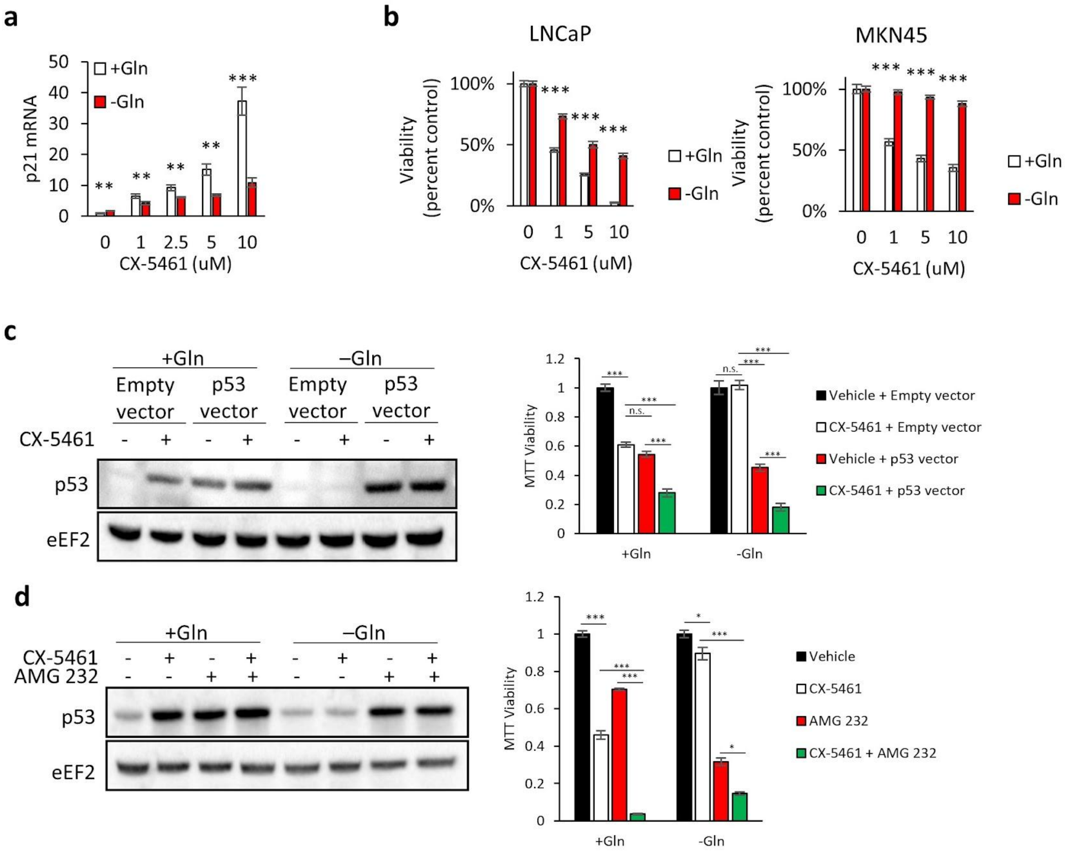


### Supplementary Fig. 6

(a) Glutamine deprivation inhibited p21 mRNA induction by CX-5461 as measured using qRT-PCR. HCT116 cells were incubated in glutamine replete (+Gln) or glutamine deprivation (-Gln) medium in the presence of vehicle (NaH2PO4) or CX-5461 at the indicated concentrations for 24 h. Data were normalized against the vehicle-treated +Gln cells.

(b) Glutamine deprivation confers cancer cells resistance to CX-5461 as assessed by MTT assay. MKN45 and LNCaP cells were incubated in +Gln or -Gln medium with vehicle (NaH2PO4) or CX-5461 at the indicated concentrations for 72 h. Data points of each treatment was normalized to the respective vehicle-treated metabolic control.

(c) p53 overexpression decreases cell viability under glutamine deprivation. HCT116 cells were transfected with empty vector (pcDNA3.1) or p53 vector. After 24 h, the transfected cells were cultured in glutamine replete (+Gln) or glutamine deprived (-Gln) media with vehicle or 1 μM CX-5461 added to the metabolic media. Cell viability was assessed 48 h later using the MTT assay.

(d) Induction of p53 with AMG 232 decreases cell viability under glutamine deprivation. HCT116 cells were cultured in glutamine replete (+Gln) or glutamine deprived (-Gln) media with a combination of 1 μM CX-5461 and 1 μM AMG 232. Cell viability was assessed 72 h later using the MTT assay.
